# Assembly and operation of a cooling stage to immobilize *C. elegans* on their culture plates

**DOI:** 10.1101/2023.02.03.526998

**Authors:** Yao L. Wang, Noa William Franklin Grooms, Claire Ma, Samuel H. Chung

**Affiliations:** Northeastern University, Department of Bioengineering, 360 Huntington Avenue, Boston, MA 02115

**Author notes:** **Corresponding author**: Samuel Chung; 360 Huntington Avenue; Boston, MA 02115. **Author contributions**: Several people contributed to the work described in this paper. SHC conceived the idea. YLW designed the stage and its assembly and operation protocols. YLW, NWFG, and CM refined the assembly and operation protocols. SHC supervised the work and the development of the manuscript. YLW wrote the first draft of the manuscript; all authors subsequently took part in the revision process and approve the final copy of the manuscript.

**Keywords:** *Caenorhabditis elegans*, animal cooling, cooling immobilization, animal processing, imaging, microscopy

## Abstract

High-resolution *in vivo* microscopy approaches can reveal subtle information and fine details inside the model animal *Caenorhabditis elegans* (*C. elegans*) but requires strong animal immobilization. Unfortunately, most current immobilization techniques require substantial manual effort, rendering high-resolution imaging low-throughput. We greatly simplify the *C. elegans* immobilization procedure by using a cooling approach that can easily immobilize entire populations of *C. elegans* directly on their cultivation plates. The cooling stage can establish and maintain a wide range of temperatures on the cultivation plate and ensures even temperature distribution across the agarose surface. In this article, we document the whole process of building the cooling stage from scratch. We envision that a typical researcher can build an operational cooling stage in their lab following our protocol without difficulty. We show how to utilize the cooling stage under three protocols, which have advantages for different experiments. We also show an example cooling profile of the stage as it approaches its final temperature and include some helpful tips in using cooling immobilization.

## Introduction

High-resolution optical microscopy provides an indispensable tool for studying in *vivo* biological structures at the subcellular level. Many biological studies require submicron resolution imaging to resolve subtle anatomical details, including neuron morphology, membrane structure, and protein localization. A high-resolution image requires an exposure time of several milliseconds to seconds, depending on the imaging modality and probe. To achieve optimal results, it is essential to carefully plan and carry out microscopy-based experiments. Crucial to this effort is an efficient animal preparation method that facilitates high-resolution imaging.

The nematode *C. elegans* is a widely utilized model organism for studying many biological processes^1^. This small animal is typically cultivated on agarose plates, and they reproduce rapidly by self-fertilization, making them well-suited for large-scale studies. Their transparency and a wide array of labeling techniques allow straightforward visualization of their internal anatomy^2,3^. The fine structures in *C. elegans* are an ideal medium for studying biological processes at the subcellular level, such as neuron regeneration^4,5^, neuron degeneration^6^, and cell division^7^. These studies necessitate imaging at submicron resolution and animal immobilization strong enough to prevent image blur. Strong immobilization is especially crucial for techniques involving multiple images in space or time, such as 3D image stacks (*i.e*., z-stacks) and timelapse. Any animal movement between the exposures can obscure the result. For *C. elegans*, strong immobilization typically involves manual manipulation of individual animals and mounting them on slides^8,9^. These time- and labor-intensive procedures make large-scale experiments very difficult. An immobilization strategy where animals are directly and reversibly immobilized on their original cultivation plates could enable high-throughput high-resolution imaging.

Cooling immobilization is an established immobilization technique for *C. elegans* and is usually combined with a microfluidic device to immobilize animals ^10–12^. However, microfluidic devices are complex, require significant operational training, and cannot be easily integrated with typical solid cultivation workflows of *C. elegans* experiments. Thus, microfluidics are not widely utilized for *C. elegans* immobilization. Here, in conjunction with our recent publication^13^, we introduce a new cooling immobilization approach using a thermoelectric cooling stage (**Figure 1**) to address these shortcomings. With the cooling stage, we can cool down a typical 60-mm polystyrene cultivation plate to any target temperature (T_set_) between −8 °C to room temperature. Our cooling stage approach can readily and reversibly immobilize an entire animal population with minimal user effort, eliminating 98% of animal processing time^13^. Below, we describe the procedures for building a cooling stage from scratch and operating it for immobilizing *C. elegans* on a typical upright microscope.

**Figure 1.**
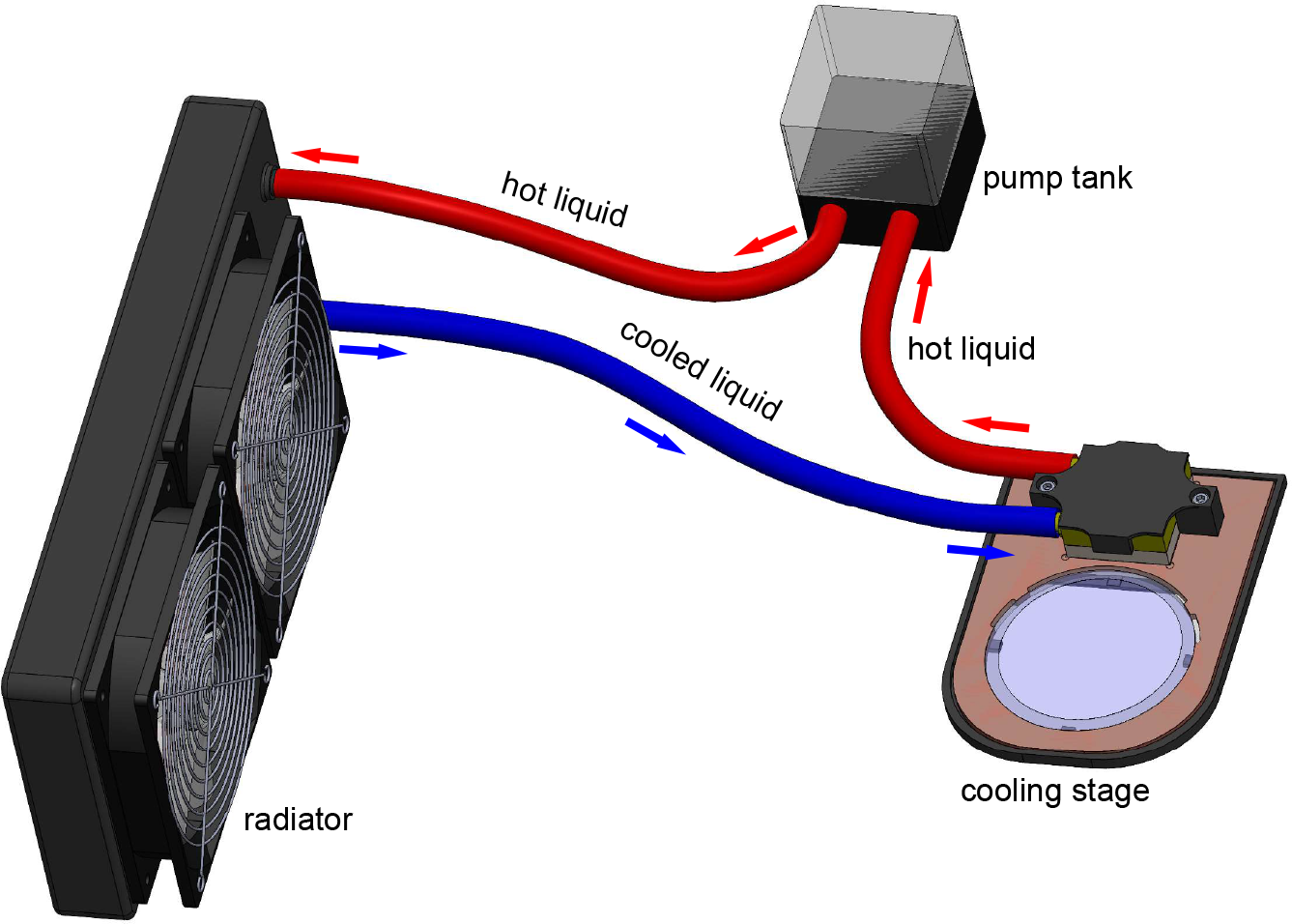
3D model of cooling stage apparatus. Electronic connections not shown for clarity. Tank pumps water through cooling block to remove heat transferred by Peltier embedded in stage. Typical 60-mm polystyrene cultivation plate sits on transparent sapphire window and is cooled by stage. Model generated in Solidworks.

## Protocol

### 1. Manufacture and prepare each component of cooling stage

Note: The cooling stage comprises several components, as listed in the **Materials List**. Most of the components are off-the-shelf. The sapphire window requires a custom order, while the copper plate, holding bracket, and isolation plate can be manufactured on-site with a computer numerical control mill or 3D printer. After the initial manufacturing, the later assembly process takes around 2-3 hours.

1.1. Use a computer numerical control mill to machine the copper plate from a 170 × 120 × 3 mm, 99.9% pure copper metal sheet (**Figure 2a**). The 2D drawing for this manufacturing is provided in the Supplementary material. Use fine-grit sandpaper to remove any sharp edges and dirty residue.
1.2. To manufacture the holding bracket and the isolation plate, use a 3D printer and 1.75mm polylactic acid (PLA) filament **(Figure 2bc)**. The 3D printer should provide at least 0.2 mm layer height for better quality.

**Figure 2.**
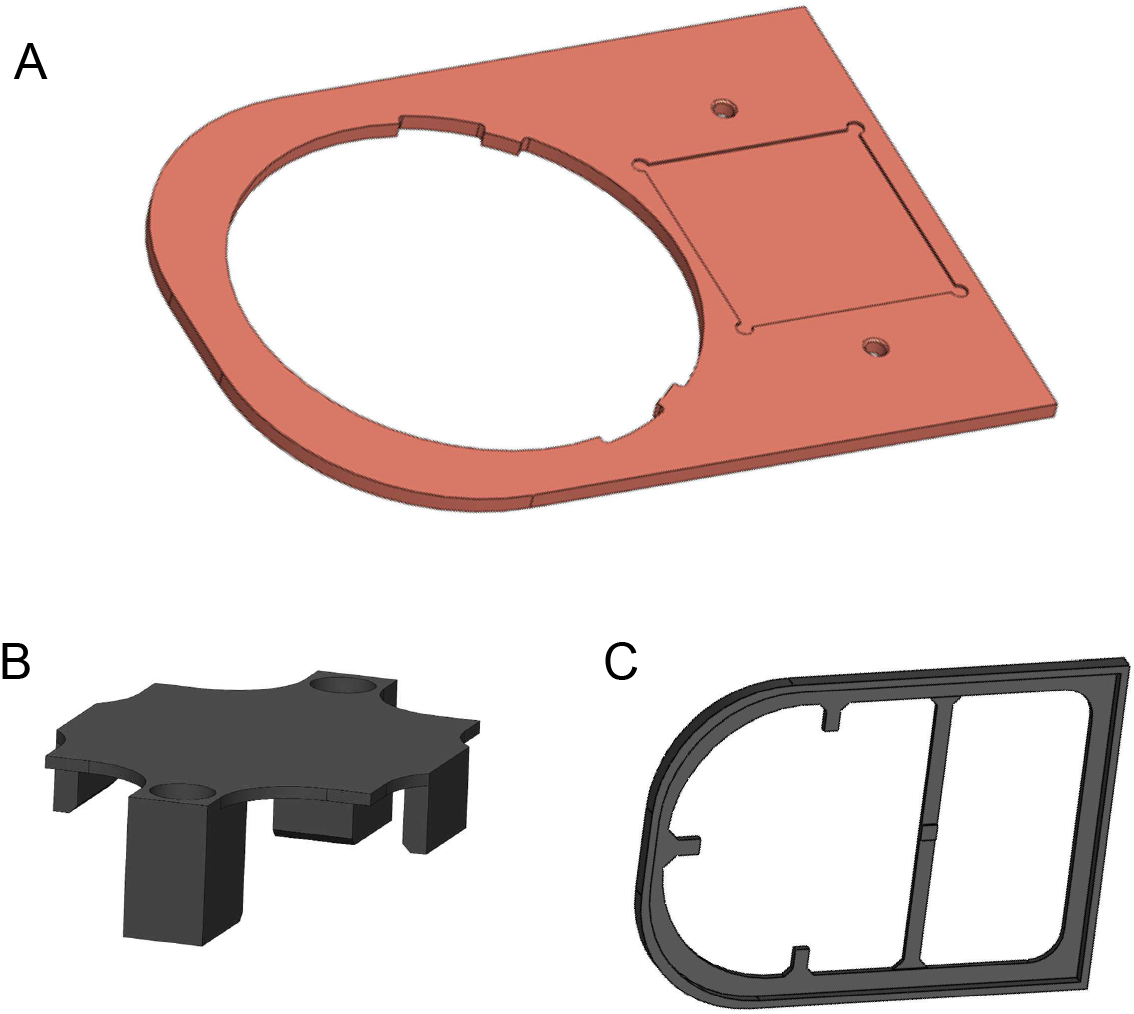
3D models of components to be manufactured. (a) Copper plate. (b) 3D-printed holding bracket. (c) 3D-printed isolation plate. Models generated in Solidworks.

### 2. Test Peltier cold and hot surfaces

Note: The Peltier, a key component of the cooling stage, is a solid-state active heat pump that transfers heat from one side to the other side^14^. One surface of the Peltier becomes hot, and the other surface becomes cold when providing electric power. Although the hot and cold sides swap if you invert the voltage input to Peltier, it is helpful to note which surface is cold when inputting standard voltages on the red (positive) and black (negative) wires. This process ensures uniformity of voltages and connections for the cooling stage.

2.1. Ensure the tunable power supply is off to prevent possible electrical hazards.
2.2. Connect the red wire of the Peltier to the positive output and the black wire to the negative output of the tunable power supply with alligator clips.
2.3. Turn on the tunable power supply, set it to around 2 V (with a sufficient setting on the current limiter), and immediately use a bare finger to feel the two surfaces of the Peltier. The cold surface becomes cold within a few seconds. After the cold surface confirmation, immediately turn off the power supply and disconnect the Peltier (**Figure 3**).
2.4. Use a marker pen to indicate the cold surface for future assembly.

**Figure 3.**
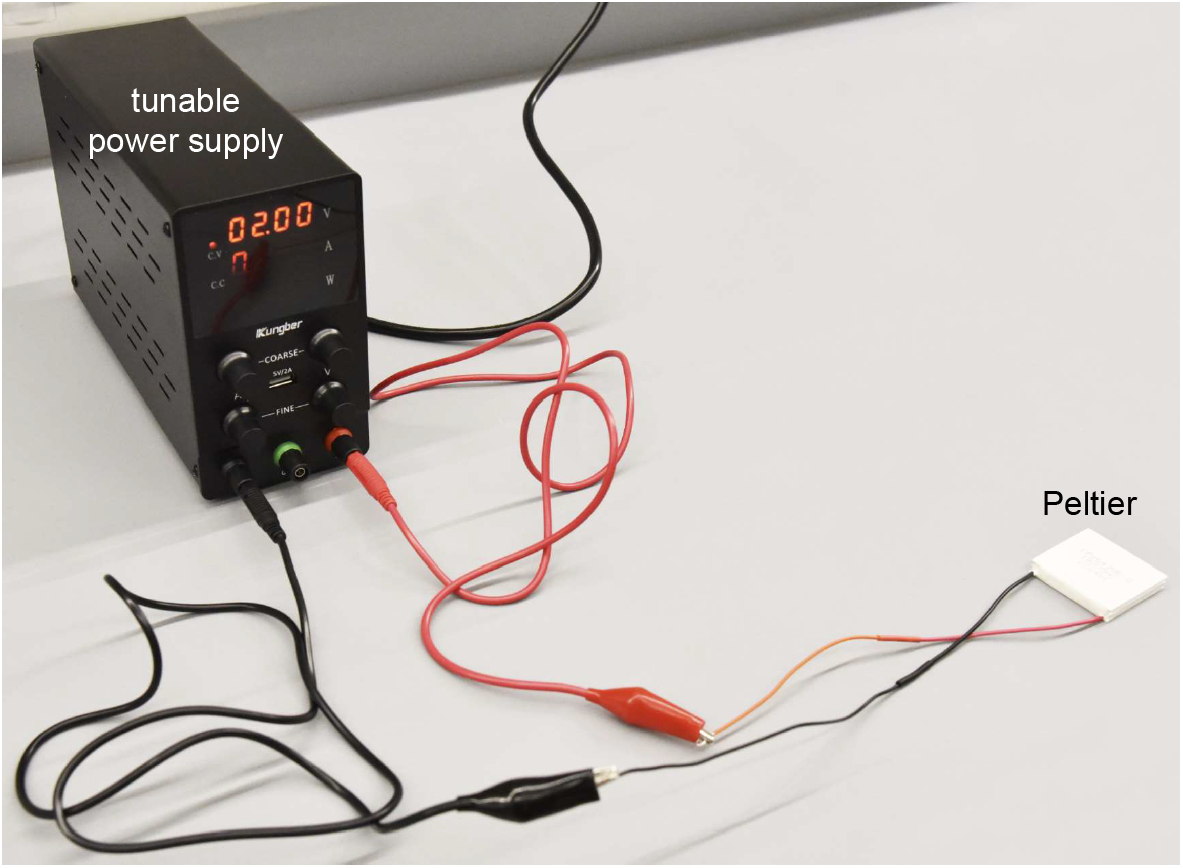
Test Peltier. Utilize tunable power supply to determine hot and cold sides of Peltier. For safety, utilize no higher than 2 V.

### 3. Constructing the water cooling assembly

Note: At this point we have every component needed for assembly. The cooling stage can be assembled with a blade, a scissor, and a hex key. Be aware of possible electric hazards due to water contact during assembling.

3.1. Prepare the Nalgene™ 50 platinum-cured silicone tube, pump tank, copper cooling block, and radiator (**Figure 4a**) for building a water cooling assembly in steps 3.2 - 3.3.
3.2. Cut the silicone tube into three sections with suggested lengths of 40 cm, 50 cm, and 80 cm. Adjust the length when needed.
3.3. Plug cut silicone tube sections into the ports of the radiator, pump tank, and copper cooling block, as shown in **Figure 4b**. Ensure all connections are water-tight. If necessary, change to a soft tube with a different diameter to ensure tightness. Do not apply paste even if the connection is not tight enough because paste may introduce clogs during future usage. Now we have the water cooling assembly, which removes heat from the cooling stage.
3.4. Prepare the water cooling assembly, a 12-V power supply, 3 red and 3 black jumper wires, a breadboard, and 500 mL water. We suggest using purified water to avoid contamination if the liquid leaks. Ensure the workbench is clear of liquid for electrical safety. Then electrically connect the water cooling assembly from steps 3.5 - 3.7.
3.5. Connect the pump tank’s and the radiator’s wires to the 12-V power supply. Ensure that the electric connections are secure and correctly made according to **Figure 4c**. Do not power up the water cooling assembly at this step. We use a breadboard for connection because of its convenience. If needed, the breadboard can be replaced with welding connections and shielding/insulation/covering for a more permanent and safe connection.
3.6. Open the pump tank cap using a flathead screwdriver. Use a funnel to add water until the pump tank is about 80% full (**Figure 4d**). Do not cap the pump tank after this filling.
3.7. Power up the water cooling assembly by plugging in the 12-V power supply or switch on (if a switch is present). After powering up, the water will begin to flow inside of the assembly, and the fans on the radiator should begin to blow. Due to the water flow from the pump tank, the liquid level in the tank will drop. Add more water to the pump tank until it stabilizes at nearly 2/3 full (**Figure 4e**). Shake the radiator to get rid of air bubbles and then cap the cooling tank. Now we have a working water cooling assembly for the cooling stage system.
3.8. Turn off the power supply before going to the next step.

**Figure 4.**
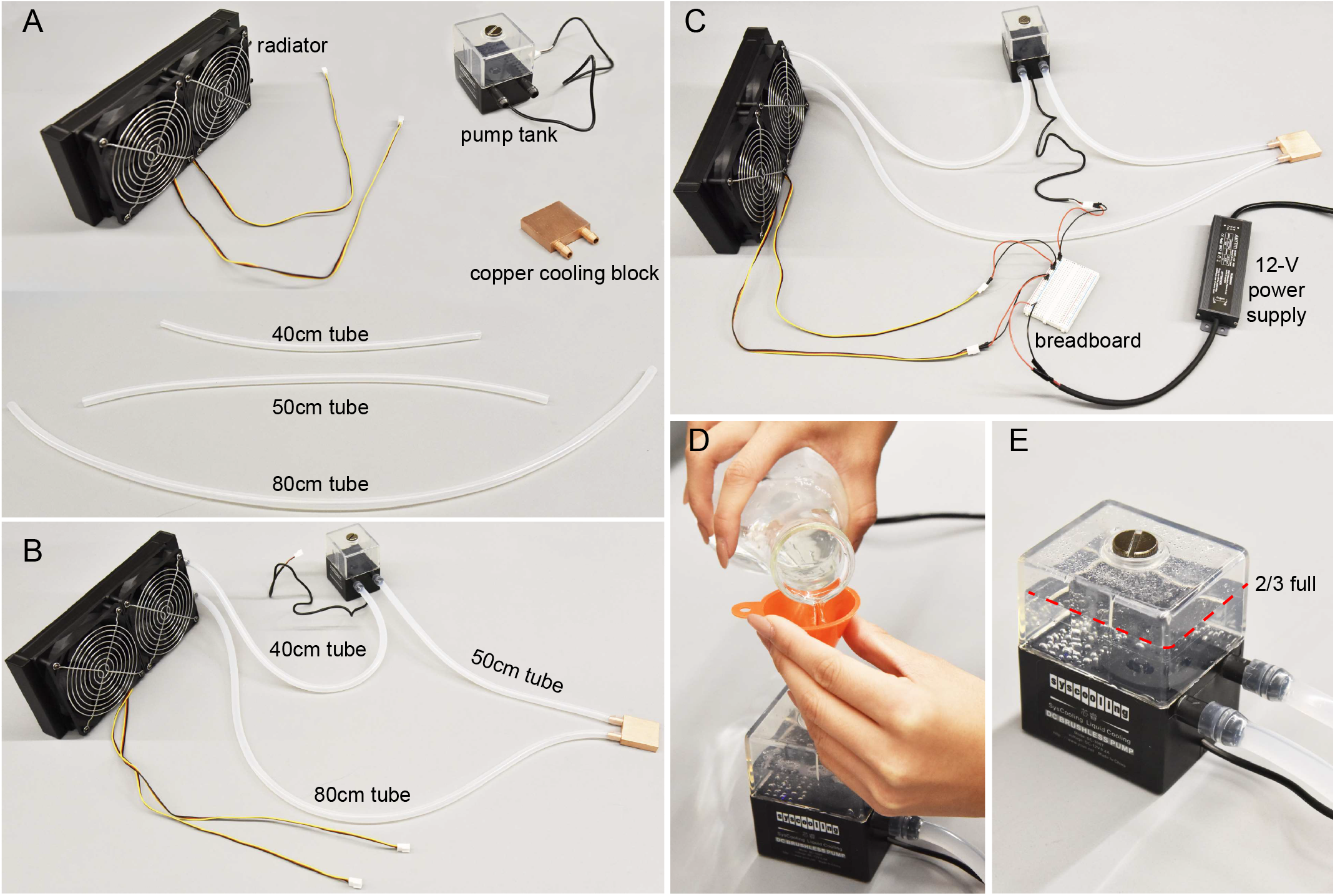
Water cooling assembly. (a) Individual components. Tubes cut to specified lengths. (b) Water cooling components connected. (c) Wires connect pump tank and radiator to 12-V power supply. In general, red (black) wires connect to positive (negative) end. (d) Purified water poured into pump. (e) Tank filled to > 2/3 full for optimal pump efficiency.

### 4. Constructing assembly to cool the Peltier using water cooling assembly

4.1. As shown in **Figure 5a**, prepare the switched-off water cooling assembly, the Peltier (cold surface marked), and the thermal paste (for improved thermal conduction). We connect the water cooling assembly to the Peltier and check the cooling capacity from steps 4.2-4.6.
4.2. Clean all surfaces of the copper cooling block with 70% ethanol (or other cleaner solution) in the water cooling assembly. Apply around 0.4 g thermal paste to one surface of the copper water cooling block. This surface should be the surface that is prefers to be facing down, based on the tube connections. Use a glove to protect your skin and try to distribute thermal paste thinly and evenly (**Figure 5b**).
4.3. Similarly, clean the hot surface of the Peltier, then apply the thermal paste to the surface (**Figure 5c**).
4.4. Connect the Peltier hot surface to the copper cooling block surface with thermal paste. Apply pressure to ensure it is secure. Note the orientation of the wires on the Peltier and the tubes of the copper cooling block, as shown in **Figure 5d**. Clean excess thermal paste.
4.5. Keep both the 12-V power supply and the tunable power supply off. Connect Peltier to the tunable power supply like in *Protocol – 2. Test Peltier cold and hot surfaces*.
4.6. Recheck both electrical and water cooling assembly connections, and then turn on the 12-V power supply and tunable power supply sequentially. Gradually turn the tunable power supply to 12 V. With the suggested Peltier, the current should be around 7.3 A. Wait for 2 mins, and the temperature of the Peltier cold surface should become colder than −35 °C. Measure this temperature with an infrared thermometer (**Figure 5e**). Do not touch the cold surface to prevent injury to hands. Check all connections and components if the temperature cannot reach below −30 °C. Air bubbles inside the water cooling assembly is a possible reason for suboptimal cooling performance.
4.7. To ensure safety in later steps, turn off the tunable power supply to Peltier, wait 1 min, then turn off the 12-V power supply to the water cooling assembly.

**Figure 5.**
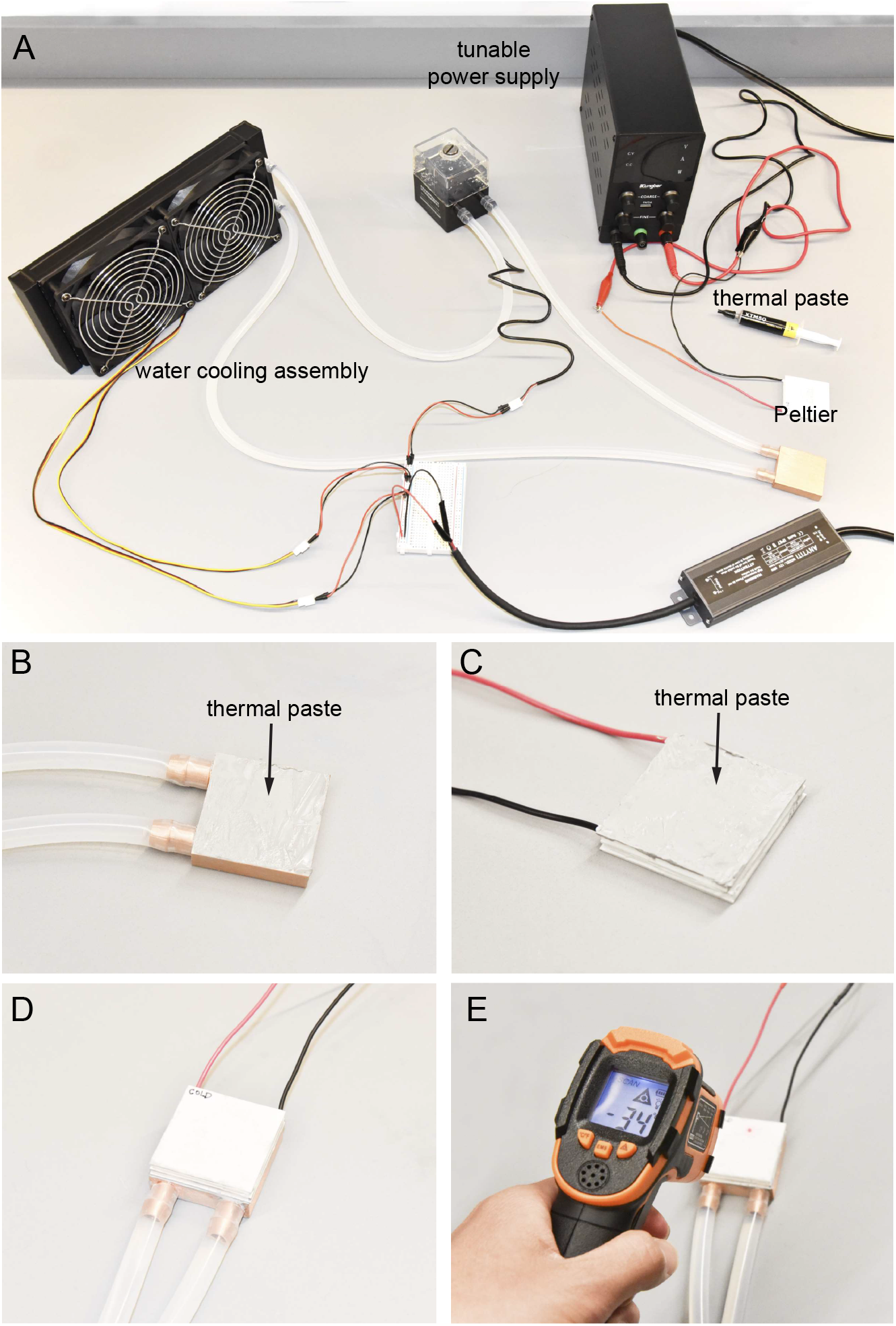
Connecting Peltier and water cooling assembly. (a) Components to operate Peltier. (b) Even application of thermal paste to surface of copper block. (c) Even application of thermal paste to Peltier hot surface. (d) Hot side of Peltier pressed onto copper block with thermal paste. (e) Infrared thermometer used to measure Peltier cold surface temperature. Ideally, the cold temperature can reach near −35 °C.

### 5. Constructing copper plate and sapphire window assembly

5.1. Prepare the copper plate, the 80-mm diameter sapphire window, thermal paste, a 4-inch-wide tape, and a sharp blade for cutting (**Figure 6a**). Carefully clean the copper plate and the sapphire window with 70% ethanol and use fine-grit sandpaper to smooth rough surfaces if necessary. The copper plate will be sealed off in the next steps, and there is no chance of cleaning it again without significant effort and reassembly. We embed the sapphire window into the copper plate from steps 5.2 – 5.6.
5.2. Apply the thermal paste on three inner surfaces, as shown in **Figure 6b**. Ensure the thermal paste covers all three surface areas but is not too thick.
5.3. Lay the copper plate on the benchtop protected with printer paper. This will make the later cleanup process significantly easier. Insert the sapphire window into the copper plate hole (**Figure 6c**). Ensure no sapphire rotation during this insertion to prevent the thermal paste from moving to other areas. Remove excess thermal paste.
5.4. Adhere the 4-inch wide tape to the top surface of the copper plate-sapphire window assembly (the surface that has the square depression area, as shown in **Figure 6d**). Avoid air bubbles between tape and copper surfaces during pasting by guiding adhesion slowly from one side to the other.
5.5. Cut out those specified blue dashed areas of the tape using a sharp blade, following **Figure 6e**. Cutting exposes the two thread holes, the square depression, and 70-mm diameter area of the sapphire window.
5.6. Tape the bottom surface of the copper plate-sapphire window assembly, and then repeat the cutting procedure (sapphire area only) on this surface, as shown in **Figure 6f**. Now, the sapphire window is fixed to the copper plate, and the copper surfaces are protected from rust.

**Figure 6.**
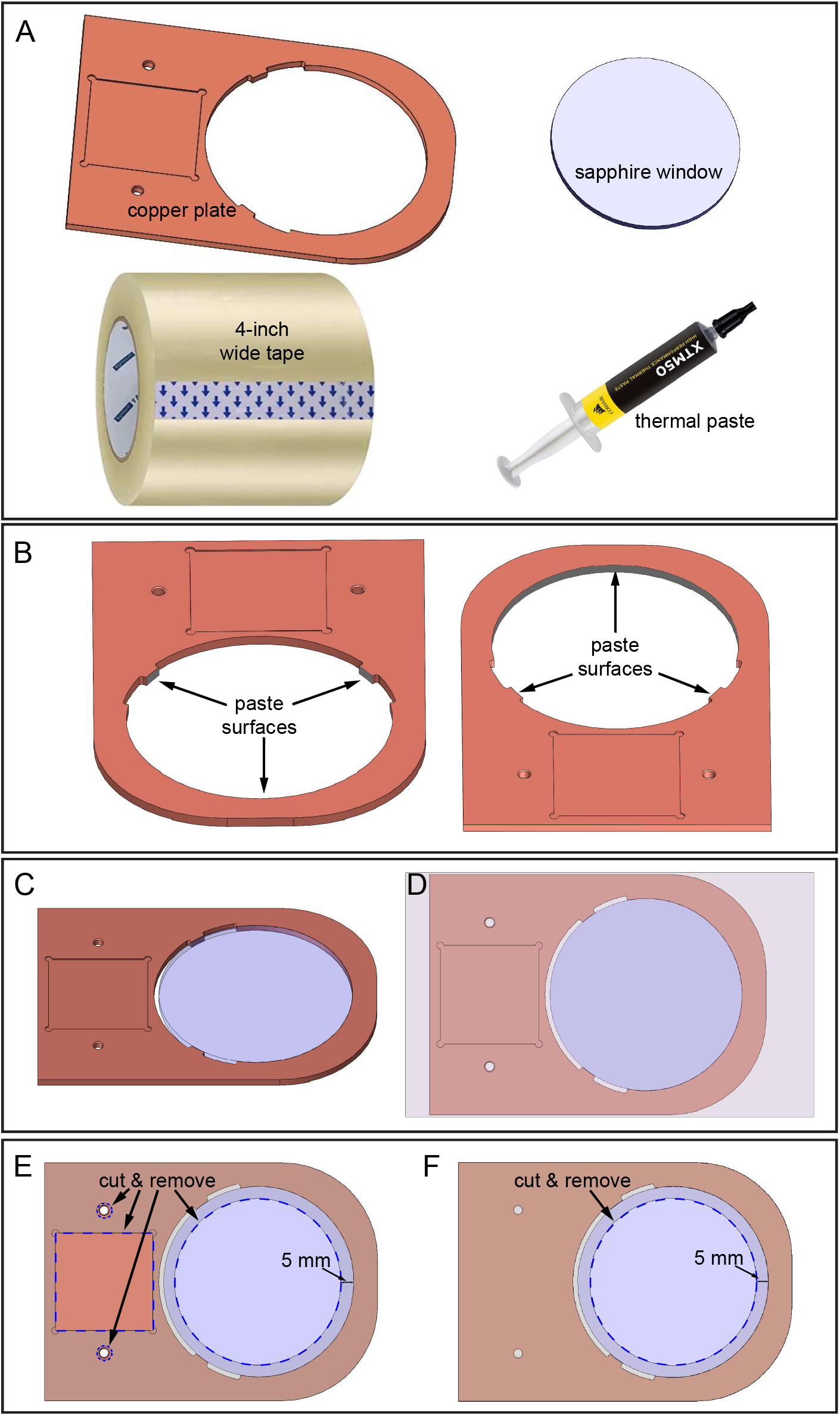
Assembling copper plate and sapphire window. (a) Components required. (b) Thermal paste applied to three inner surfaces of copper plate where the sapphire window will contact. Two downward-looking views of copper plate showing location of three surfaces. (c) Sapphire window in copper plate hole. (d) Tape applied to top surface of assembly. (e) Top-side: Blue dashed lines indicate locations to cut and remove tape: square depression, two holes, and a 70-mm diameter sapphire area. (f) Bottom-side: Cut and remove the tape as shown.

### 6. Cooling stage final assembling

6.1. All essential sub-assembly and components are ready now. We can begin the final assembling from step 6.2 – 6.5.
6.2. Apply about 0.4 g thermal paste to the square depression of the copper plate (**Figure 7a**).
6.3. Apply about 0.4 g thermal paste to the cold surface of the Peltier. Note that the Peltier is already attached to the copper cooling block (**Figure 7b**).
6.4. Connect the Peltier cold surface to the copper plate depression with downward pressure. Clean all excess thermal paste (**Figure 7c**).
6.5. Mount the 3D-printed bracket on the top of the copper cooling block, and then use a hex key to tighten two 8-32, 1/2 long screws to fix the bracket to the copper plate (**Figure 7d**). Only low-torque tightening is recommended as the 3D printed bracket may break. This will ensure proper thermal conduction from the Peltier to the copper.
6.6. Place the copper plate into the 3D-printed isolation base for thermal isolation from the benchtop or microscope base during operation (**Figure 7d**).
6.7. The cooling stage is assembled and ready to use (**Figure 7e**).
6.8. For microscopy, place the completed cooling stage on an upright microscope platform (**Figure 8a**).

**Figure 7.**
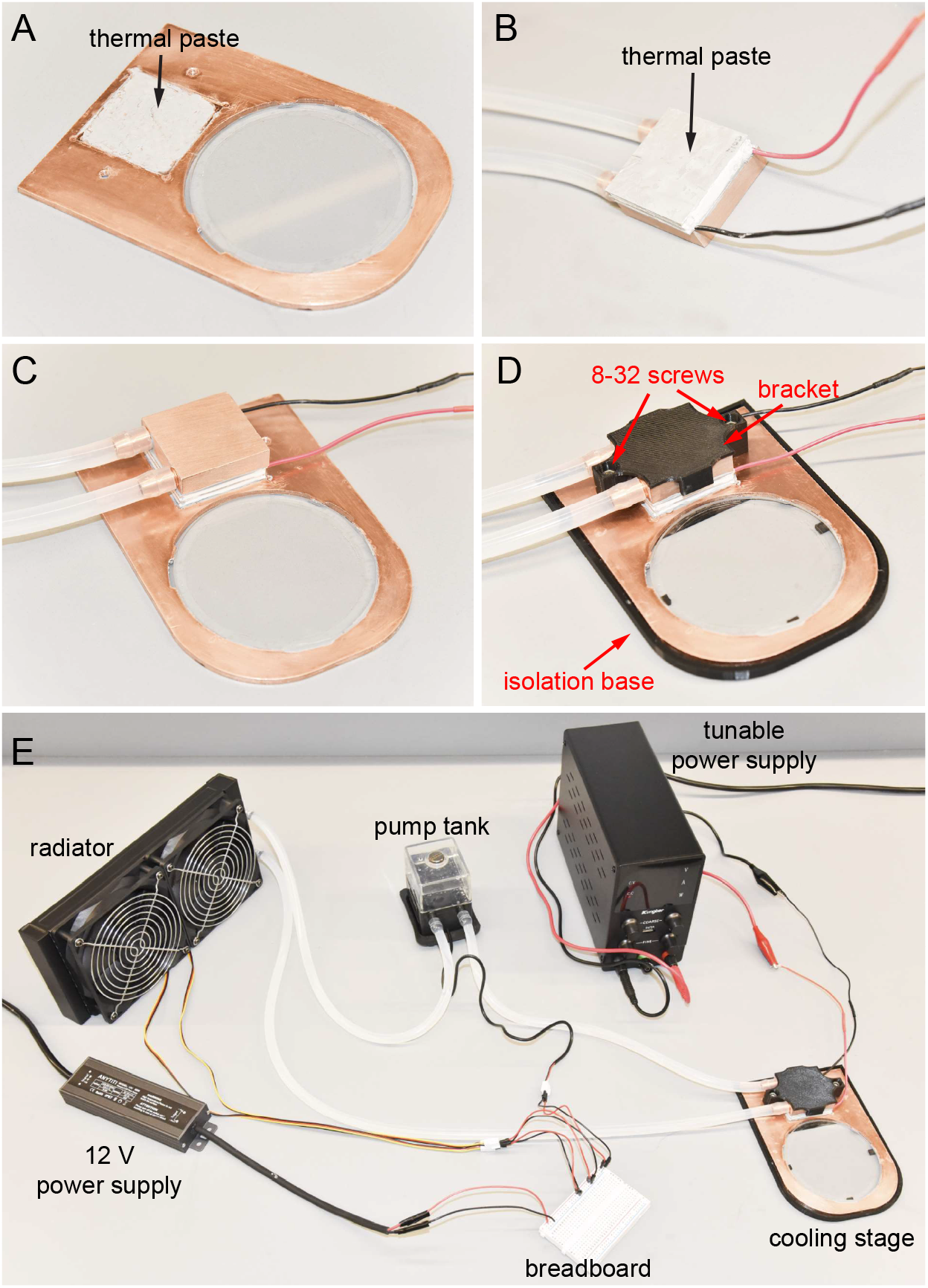
Cooling stage final assembly. (a) Thermal paste applied to depression of copper plate. (b) Thermal paste applied to cold side of Peltier. (c) Cold surface of Peltier connected to depression. (d) Copper cooling block fixed to copper plate using screws. Cooling stage in isolation base. (e) Completed cooling stage assembly.

**Figure 8.**
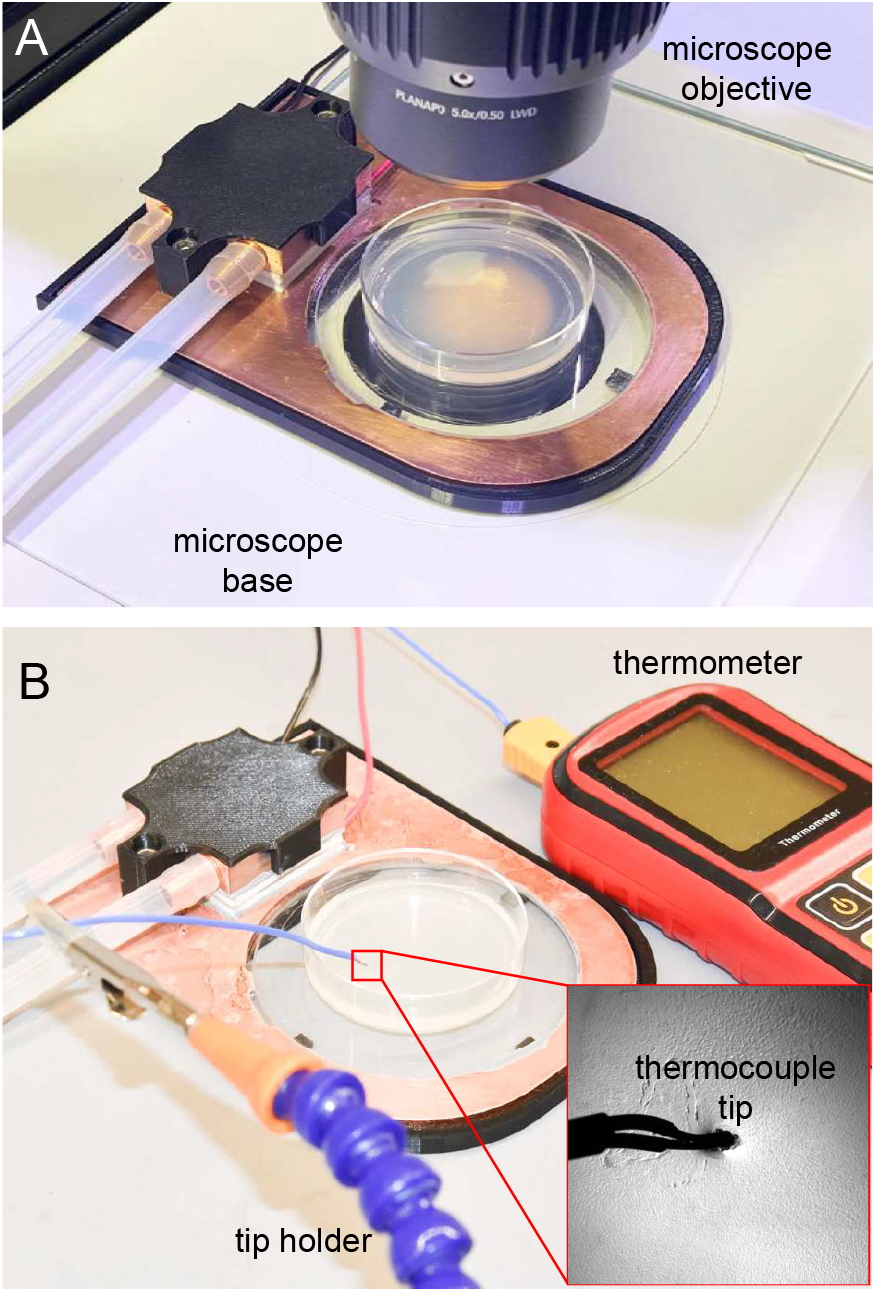
Cooling stage on microscope and thermocouple measurement. (a) Cooling stage placed on microscope base for imaging. Sapphire window is transparent, allowing transillumination. (b) Thermocouple thermometer used to measure agar surface temperature. Tip inserted around 1mm into the agar.

Note: We finished the assembling of the cooling stage. Next, we further introduce three different cooling strategies with different cooling rates and user effort (compared in **Table 1**). The preferred strategy may depend on application. We show protocols for those three cooling immobilization strategies, which we term “slow”, “fast”, and “abrupt”. Further details available from our companion publication, which fully characterized the strategies and animal movement^13^.

**Table 1.** Cooling strategies comparison

|  | slow cooling | fast cooling | abrupt cooling |
| --- | --- | --- | --- |
| stage occupation | minimum | long | medium |
| time until animals are immobilized | long | medium | very short |
| immobilization strength | strong | medium | medium |
| user effort | minimum | slightly more than minimum | maximum |

Note: In a real experiment, the parameters provided in this manuscript, such as voltages and times, will depend on specific properties of the cultivation plates and stage, such as the amount of agar in the plates, the stage’s efficiency, and the ambient temperature and humidity.

Note: Different temperatures can immobilize animals differently due to animals cultivation temperature, strain mutations, and many other conditions.

Note: For initial experiments, we suggest keeping track of the agar surface temperature to ensure animals are properly immobilized at 6 °C. For temperature measurement, sterilize the thermocouple tip of the thermometer (Proster, San Jose, CA) using 70% ethanol solution and insert the tip 1 mm into the agarose to ensure an accurate temperature reading. Use a clamp holder or other holders to hold the thermometer tip (**Figure 8b**). Future experiments that are largely replicated from the initial one can utilize the same parameters, usually without frequent temperature tracking.

### 7. Slow cooling immobilization protocol

Note: Slow cooling strategy is primarily useful for immobilizing 20 °C cultivated N2 animals at 6 °C. 15 °C-cultivated N2 animals are most immobilized at 1 °C.

7.1. Place a lidded cultivation plate with animals to a 4 °C refrigerator for 1 hour.
7.2. After moving the cultivation plate to the refrigerator, turn on the cooling stage’s 12-V power supply and turn the tunable power supply voltage to 5.5 V.
7.3. After the lidded cultivation plate has stayed in the 4 °C refrigerator for 1 hour, transfer the plate immediately to the cooling stage and remove the lid (**Figure 8a**). Such plates are usually around 6 °C. The pre-cooled stage is stable and cold enough to maintain the agar surface at 6 °C. If the agar surface temperature changes, as measured or by noting animal movement, slightly adjust the voltage until it stabilizes at 6 °C.
7.4. Animals are properly immobilized at the time of transfer.

### 8. Fast cooling immobilization protocol

Note: Fast cooling strategy is the most basic immobilization method; however, agar plates idly occupy the stage for an extended time while reaching the T_set_. Also, when strong immobilization is needed and the T_set_ is 6 °C, idle time is extended to around 1 hour^13^.

8.1. Turn on the cooling stage’s 12-V power supply and set the tunable power supply voltage to around 12-V. Wait for 10 mins.
8.2. Bring a cultivation plate from its incubator directly to the cooling stage and remove the lid. The plate begins to cool.
8.3. Once the agar surface temperature decreases to (T_set_ + ΔT) °C, adjust the tunable power supply to V_set_ and wait until the agar reaches T_set_. The V_set_ is the appropriate voltage to stabilize the agar at T_set_. The ΔT is a variable that prevents overcooling. See **Table 2** for the combination of T_set_,, ΔT, and V_set._ Note that the data in this **Table 2** is based on our environment and usage conditions, and it should be revised accordingly in use.
8.4. Animals are immobilized when the agar reaches T_set_. Immobilization improves with time until ~50 min after initiation of cooling.

**Table 2.** Parameters to achieve desired temperature in fast cooling strategy

| $T_{\text{set}}$ (°C) | $\Delta T$ (°C) | $V_{\text{set}}$ (V) |
| --- | --- | --- |
| 1 | 2 | 8 |
| 2 | 3 | 7.4 |
| 3 | 4.5 | 7 |
| 4 | 5.5 | 6.5 |
| 5 | 6 | 5.9 |
| 6 | 6 | 5.5 |

### 9. Abrupt cooling immobilization protocol

Note: Abrupt cooling strategy consumes the most user time but immobilizes animals the most rapidly from their cultivation temperature.

9.1. Turn on the cooling stage’s 12-V power supply and turn the tunable power supply voltage to around 12-V. Keep for 10 mins.
9.2. Bring an unoccupied agar plate to the cooling stage. Use Step 8.3 in the fast cooling immobilization protocol to stabilize the agar surface temperature at T_set_.
9.3. Move animals from their original cultivation plate to the cooled plate sitting on the cooling stage.
9.4. Based on the small animal size, we expect animals are cooled to T_set_ in seconds and immobilized. Immobilization improves with time until ~50 min after initiation of cooling.

### 10. Animals revive after cooling immobilization

10.1. Worms can revive to their normal status after moving the cooled culture plate back to their original incubator or room temperature. It usually takes 20 mins to 1 hour until all worms on that plate revive to their normal crawling and feeding behavior.

## Results

### Temperature measurement with an infrared camera

We designed the cooling stage to ensure the temperature distribution in the plate’s central 40 mm diameter area is uniform. We use a FLIR infrared camera to image the temperature distribution on the agar surface. The maximum temperature difference is around 1 °C when the T_set_ is 1 °C, 3 °C or 6 °C (**Figure 9a**).

**Figure 9.**
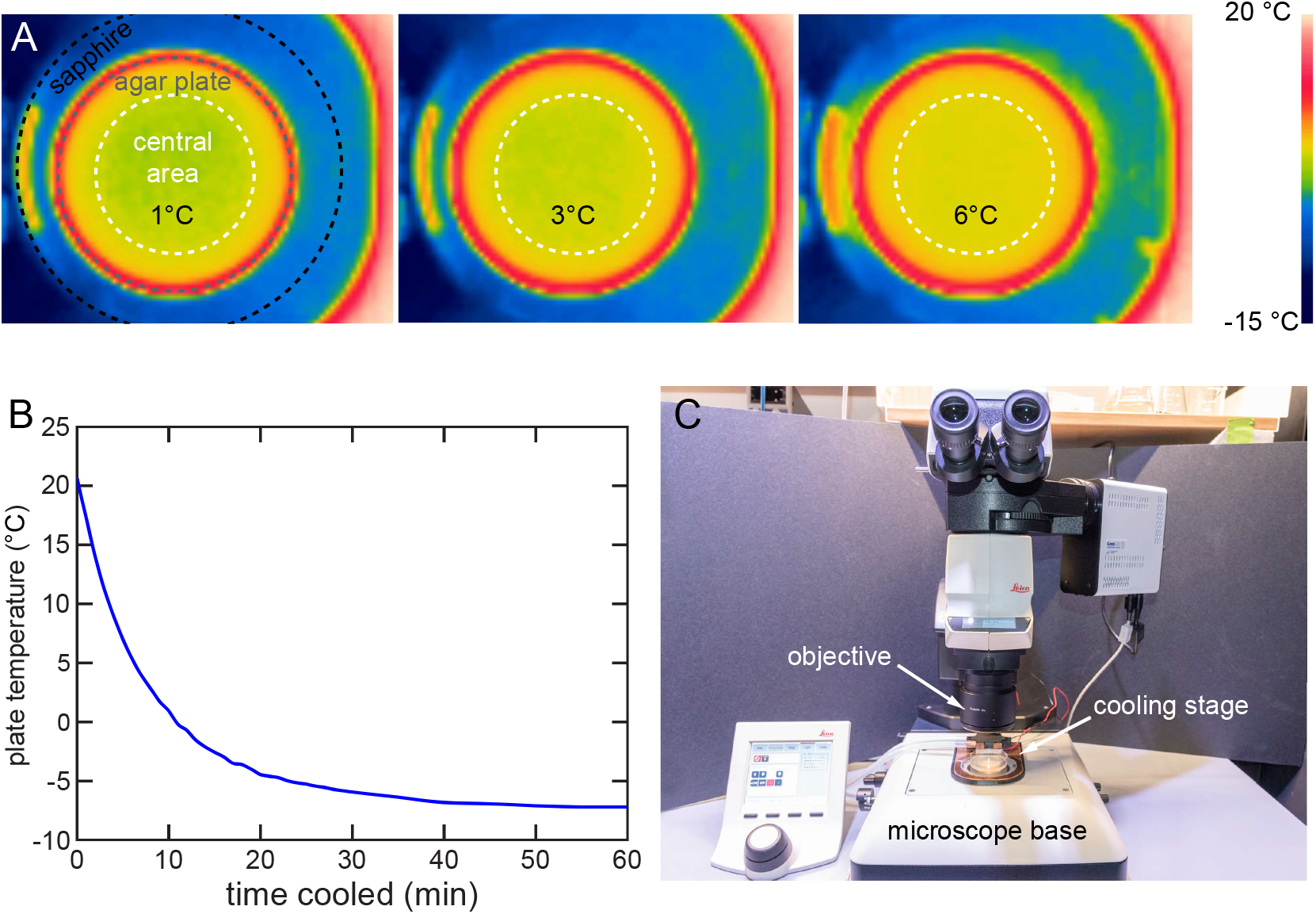
Cooling stage characterization and usage. (a) Thermal images showing agar surface cooled to 1, 3, and 6 °C. Even temperature distribution within central 40-mm area (white dashed circle). (b) Temperature of agar surface over time on cooling stage at 12 V. Agar surface can be cooled below −7 °C. Temperature measured by method in Fig. 8a. (c) Cooling stage in use on typical upright microscope. Cooling stage can be easily installed or removed.

### Cooling rate with the fast cooling approach

We used the fast cooling strategy to characterize the cooling rate of a stage at 12-V. We placed a 20 °C plate on the cooling stage and used the thermocouple thermometer to track the surface temperature. The stage cools down the 20 °C plates to 6 °C in 6 mins, to 1 °C in 10 mins, and eventually stabilizes below −7 °C in around 40 min (**Figure 9b**).

### Use the cooling stage on an upright microscope platform

An upright microscope typically comprises an objective for imaging, a stage for sample holding, and illumination. Our cooling stage is designed for use on a typical upright microscope stage with easy insertion and removal (**Figure 9c**). When cooling immobilization is needed for imaging or screening, simply place the cooling stage on the microscope stage to finish the installment and vice versa.

### Helpful tips for using the cooling stage

Based on experience, we provide some notes for device operation to ensure an efficient cooling immobilization experiment.

1. Check whether there are any air bubbles inside the water cooling assembly. Air bubbles degrade the cooling to the Peltier hot surface and thus degrade the cooling effectiveness of the cooling stage. If air bubbles are present, turn on the 12-V power supply to make the water flow and shake all the components of the water flow. Air bubbles can be flushed out from trapped areas and be vented by the pump tank.
2. Ensure that the water flow tubing is not bent or crossed in any way when assembling the water cooling assembly. Tube bending or crossing may prevent adequate flow of water and reduce cooling efficacy.
3. Room humidity affects the cooling performance and introduces condensation and ice on the cooling stage. Before placing a cultivation plate on the cooling stage, it is recommended to use a paper tissue to remove condensation or use a heatsink to remove ice formed on the sapphire window quickly.
4. The pump tank and radiator fans can cause small vibrations to the microscope if they work on the same table. Microscope vibration blurs the image acquired and should be avoided. Use a cushion to mechanically insulate the tank and radiator, or place them on a separate nearby table.
5. The cooling stage can become a heating stage by reversing the electrical connection to the Peltier.

## Discussion

We show the cooling stage manufacturing, assembling, and usage in this manuscript. Most of the components are off-shelf items that can be purchased online. Some components, like the copper plate and the sapphire window, need custom order and may take up to a month to fabricate. Other components that can be 3D printed are easily fabricated in most research institutions. The assembling process needs only a few tools and can be quickly done by a non-expert in a few hours. Thus, we believe that most biological laboratories can easily deploy this device.

The cooling stage is a low-cost, highly effective immobilization device for *C. elegans*. By fine-tuning the temperature on animals, we conclude that cooling immobilization can be nearly as strong as conventional sodium azide immobilization. Cooling is appropriate for high-resolution imaging, screening, or any other research requiring strong animal immobilization.

There are several limitations of this cooling stage immobilization. First, when cooled to the same temperature, animals raised at different temperatures are immobilized to different degrees. Users should fine-tune parameters for any experiment, but particularly for animals that were not cultivated at 20 °C or have temperature-sensitive mutations. Second, the cooling stage is not designed for an inverted microscope. However, by slight modification, we envision it can be used on an inverted microscope with the same overall concept. Third, imaging or screening on a cultivation plate directly may introduce contamination to the plate, especially because we prefer to keep the lid off for best imaging quality in high resolution imaging. We encourage users to complete their experiments expeditiously to minimize contamination.

## Supporting information

Materials List

## Acknowledgements

We acknowledge Noah Joseph (Northeastern Bioengineering Dept.) for copper plate machining.

