## Supplementary material for "Assembly and operation of a cooling stage to immobilize *C. elegans* on their culture plates": Copper plate in Metric.pdf

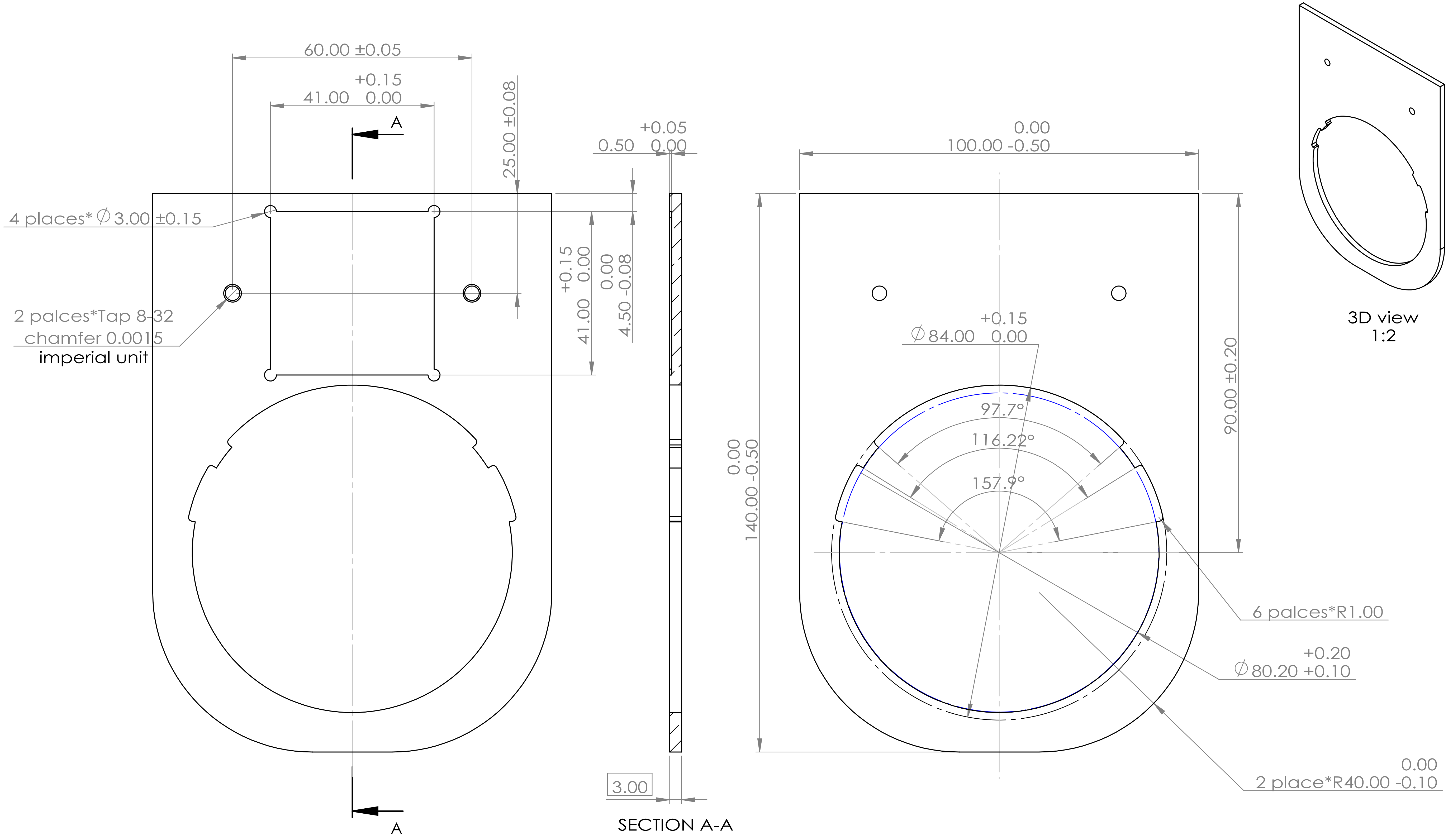


1. All unit are in mm or degree, except the 8-32 thread Tap;
2. Chip removing on every outer edge.

| | | | | |
|---|---|---|---|---|
| UNLESS OTHERWISE SPECIFIED: DIMENSIONS ARE IN MILLIMETERS SURFACE FINISH: TOLERANCES: LINEAR: ANGULAR: | FINISH: | DEBURR AND BREAK SHARP EDGES | DO NOT SCALE DRAWING | REVISION |
| | NAME | SIGNATURE | DATE | TITLE: Copper plate in Metric |
| DRAWN | | | | |
| CHK'D | | | | |
| APPV'D | | | | |
| MFG | | | | |
| Q.A | | | MATERIAL: | DWG NO. A3 |
| | | | WEIGHT: | SCALE:1:1 SHEET 1 OF 1 |

RK
